# Machine Learning Workflow for Single-Cell Antimicrobial Susceptibility Testing of *Klebsiella pneumoniae* to Meropenem in Sub-Doubling Time

**DOI:** 10.1101/2022.11.03.515093

**Authors:** Kristel C. Tjandra, Nikhil Ram-Mohan, Manuel Roshardt, Elizabeth Zudock, Zhaonan Qu, Kathleen E. Mach, Okyaz Eminaga, Joseph C. Liao, Samuel Yang, Pak Kin Wong

**Author notes:** These authors contributed equally to this work (Kristel C. Tjandra, Nikhil Ram-Mohan, Manuel Roshardt).

## Abstract

Multidrug-resistant *Enterobacteriaceae* are among the most urgent global public health threats associated with various life-threatening infections. In the absence of a rapid method to identify antimicrobial susceptibility, empirical use of broad-spectrum antimicrobials such as carbapenem monotherapy has led to the spread of resistant organisms. Rapid determination of antimicrobial resistance is urgently needed to overcome this issue. By capturing dynamic single-cell morphological features of over thirty-nine thousand cells from nineteen strains of *Klebsiella pneumoniae*, we evaluated strategies based on time and concentration differentials for classifying its susceptibility to a commonly used carbapenem, meropenem, and predicting their minimum inhibitory concentrations (MIC). We report morphometric antimicrobial susceptibility testing (MorphoAST), an image-based machine learning workflow, for rapid determination of antimicrobial susceptibility by single-cell morphological analysis within sub-doubling time. We demonstrated that our algorithm has the ability to predict MIC/antimicrobial susceptibility in a fraction of the bacterial doubling time (<50 min.). The classifiers achieved as high as 97% accuracy in 20 minutes (two-fifths of the doubling time) and reached over 99% accuracy within 50 minutes (one doubling time) in predicting the antimicrobial response. A regression model based on the concentration differential of individual cells from nineteen strains predicted the MIC with 100% categorical agreement and essential agreement for seven unseen strains, including two clinical samples from patients with urinary tract infections with different responsiveness to meropenem. The expansion of this innovation to other drug-bug combinations could have significant implications for future development of rapid antimicrobial susceptibility testing.

## Introduction

The emergence of multidrug-resistant pathogens is a worldwide calamity. Multidrug-resistant bacteria, such as the carbapenemase-producing *Enterobacteriaceae*, pose an increasing threat to public health due to their high mortality rate and rapid acquisition of resistance to available antimicrobials.^1,2^ *Enterobacteriaceae* are common causes of nosocomial infections and could lead to various life-threatening infections, including severe lung, urinary tract, and bloodstream infections. Broad-spectrum carbapenems like meropenem, an intravenous beta-lactam antimicrobial, are often adopted for *Enterobacteriaceae* nosocomial infections until the antimicrobial susceptibility test (AST) results are available to guide more targeted therapy. In the absence of a rapid method to determine antimicrobial susceptibility, empirical use of antimicrobials is warranted as urgent treatment is linked with improved outcomes. Inappropriate antimicrobial treatment results in almost twice higher mortality rate in infected patients and accelerates the emergence and spread of superbugs.^3^ Therefore, rapid determination of antimicrobial susceptibility to guide treatment is crucial to save lives and curb the widespread of multidrug-resistant pathogens.

Phenotypic AST, such as broth microdilution and Kirby-Bauer disk diffusion test, evaluates the ability of an antimicrobial to inhibit bacteria growth and is the gold standard for determining antimicrobial susceptibility. The turbidity of liquid media or the formation of bacterial colonies provides a measure of bacteria growth with the presence of an antimicrobial and generates quantitative minimum inhibitory concentration (MIC).^4^ Nevertheless, phenotypic AST, which relies on bacterial growth for 18 hours or more, is incompatible with a rapid turnaround. This limitation of current AST propelled the development of novel approaches to improve both speed and accuracy.^5–9^ Single-cell imaging analysis is an emerging strategy that offers the possibility of reducing the turnaround time and improving the diagnostic resolution. By visualizing the replication of individual cells, the response of bacteria to antimicrobials can be in principle reduced to one or a few doubling times of the bacteria.^8–12^ However, rapid AST techniques that rely on the area occupied by the cells as a measure of the growth could mistake a transient increase in cell sizes due to antimicrobial tolerance as growth.^9^ Phenotypic variants, drug accumulation, and growth phase can also introduce uncertainties in rapid AST assays.^13,14^ Therefore, existing approaches relying on cell replication often require at least 2 hours, if not more, to deliver reliable results, especially for slow-growing and fastidious bacteria.^9,15–18^ While there are ample examples of rapid AST, the search for an assay that can deliver AST results (1) rapidly in a point-of-care timeframe, (2) quantitatively with MIC determination, and (3) efficiently with a small inoculum size is still ongoing.

A promising strategy for rapidly determining bacteria response to antimicrobials is to monitor their morphological changes.^19^ Bacteria undergo a wide variety of morphological changes in response to the environment.^20^ These changes, such as filamentation, bulging, and lysis, indicate stress in bacteria and have been applied for investigating the mechanisms of action of antimicrobials.^7,21–23^ For instance, distinct morphological transformations could be induced in *Escherichia coli* depending on the type of beta-lactam antimicrobials.^24^ *Pseudomonas aeruginosa,* which are known to be highly tolerant against beta-lactams, undergo a transition from rod-shaped to viable spherical cells when treated with meropenem.^25^ The minute change in the bacterial morphology could be indicative of the bacterial response to antimicrobials before cell replication. While promising examples of bacterial analysis at the single-cell level have emerged,^26–28^ the potential of single-cell morphological analysis has not been realized for rapid AST and MIC determination due to a lack of effective strategy for analyzing the dynamic, multiparametric morphological features of bacteria.

Herein, we demonstrate a machine learning workflow, termed morphometric antimicrobial susceptibility test (MorphoAST), for single-cell *Klebsiella pneumoniae* AST and MIC quantification against meropenem in a fraction of the bacterial doubling time. The workflow combines single-cell imaging, computer vision feature extraction, and supervised learning models for predicting the response of bacteria to antimicrobials. We measured dynamic morphological features of individual *Klebsiella pneumoniae* in the presence of meropenem. We processed the data using time and concentration differential strategies and trained machine learning models to predict the antimicrobial response and MIC. The models were cross-validated and tested against cells from seven unseen strains, including two samples from patients with urinary tract infections to provide an unbiased evaluation of the trained model. The results were reported according to the CLSI performance standards for antimicrobial susceptibility testing (M100) guideline.

## Results

### The MorphoAST workflow for rapid *Klebsiella pneumoniae* AST

We developed a machine learning workflow for rapid AST determination for *K. pneumoniae* against meropenem in a sub-doubling time of the bacteria (Figure 1A). The MorphoAST workflow started with imaging of individual bacteria under various antimicrobial concentrations. Cells were grown to a log phase and treated with varying concentrations of meropenem. Cells without any antimicrobials were imaged in the same manner as controls for the experiment. Live bacteria were mounted on an agarose pad to minimize cell movement due to cell motility and Brownian motion. Time-lapse images were taken every 5 minutes. Images were aligned to match the location of individual cells at each frame and then subject to MicrobeJ analysis for extracting morphological features.^29^) A total of 21 automatically-extracted parameters (Supplementary Table 2) describing the cellular dimension and orientation were generated. The data have been analyzed either fitting with an exponential function to extract the feature changing rates (Figure 1B) or normalized against the untreated control to extract the feature differentials (Figure 1C). The data were applied to train and validate machine learning classifiers to predict the antimicrobial response (Figure 1D) and regressors to predict the MIC of the bacterial strain against the antimicrobial (Figure 1E).

**Figure 1.**
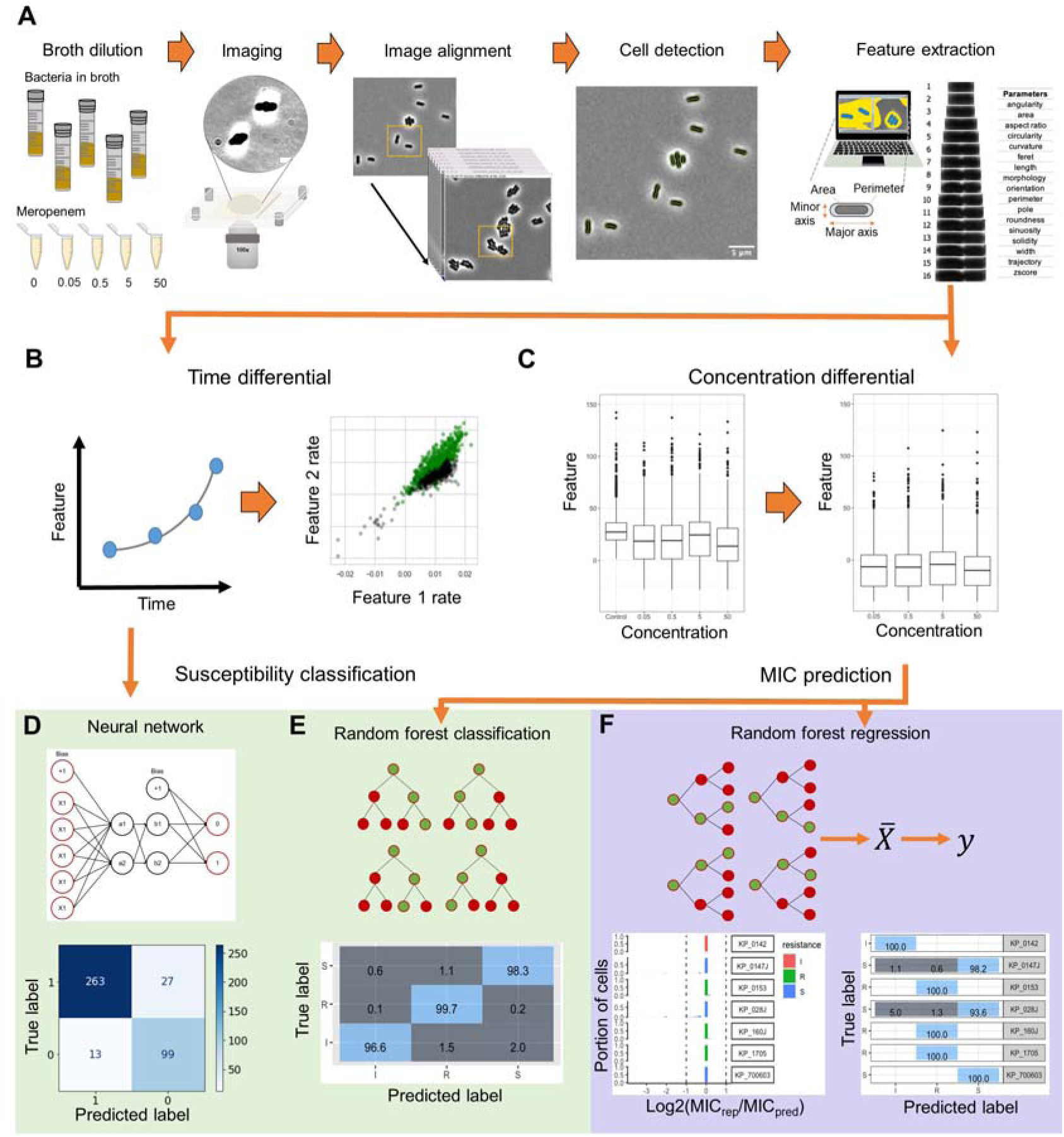
A schematic flowchart of the single-cell MorphoAST workflow for susceptibility classification and minimum inhibitory concentration (MIC) prediction. (A) The workflow starts with time-lapse imaging of individual bacteria under various antimicrobial concentrations. The morphological features of the bacteria are extracted automatically using MicrobeJ, an ImageJ plugin. (B) In the time differential approach, the feature changing rates are extracted by exponential curve fitting from the time-lapse data. (C) In the concentration differential approach, all features extracted from MicrobeJ are normalized against the feature mean of the control population. All datasets are visualized and separated into training and validation sets. (D-E) The feature-changing rate datasets are applied to train an artificial neural network to create a classification model. In parallel, the normalized dataset is trained using a random forest classifier with cross-validation. The trained models are then validated with the validation dataset for analyzing the prediction accuracy for susceptibility classification. (F) A Random Forest regression model is generated by the normalized data. The regressor model is validated through cross-validation and tested against unseen cells for evaluating categorical agreement and essential agreement of the assay.

### Single-cell imaging and feature extraction

A total of 24 *K. pneumoniae* strains bearing various carbapenem resistance genes and sensitivity towards meropenem were monitored over time (Supplementary Table 1). Changes in bacterial morphology were observed over 90 minutes post-antimicrobial incubation. Only the first 50 minutes, which is approximately the doubling time of *K. pneumoniae*, was analyzed. Each strain showed a unique morphological response to the varying antimicrobial concentration based on their MIC. Figure 2A and B show *K. pneumoniae* (KP0142) treated with and without meropenem. Bulging of the bacteria was only observed in the meropenem case. Supplementary Figure 1 compares three strains (KP0016 – Susceptible, KP0142 – Intermediate, and KP0143 – Resistant) at the early time points (see also movies S1-3). Similarly, at 5 μg/mL meropenem, which is higher than the breakpoint MIC for meropenem, susceptible and intermediate strains (KP0016 and KP0142) showed noticeable ‘bulging’ or protrusion around the center of the cell that was not observed in the resistant strain (KP0143).

**Figure 2.**
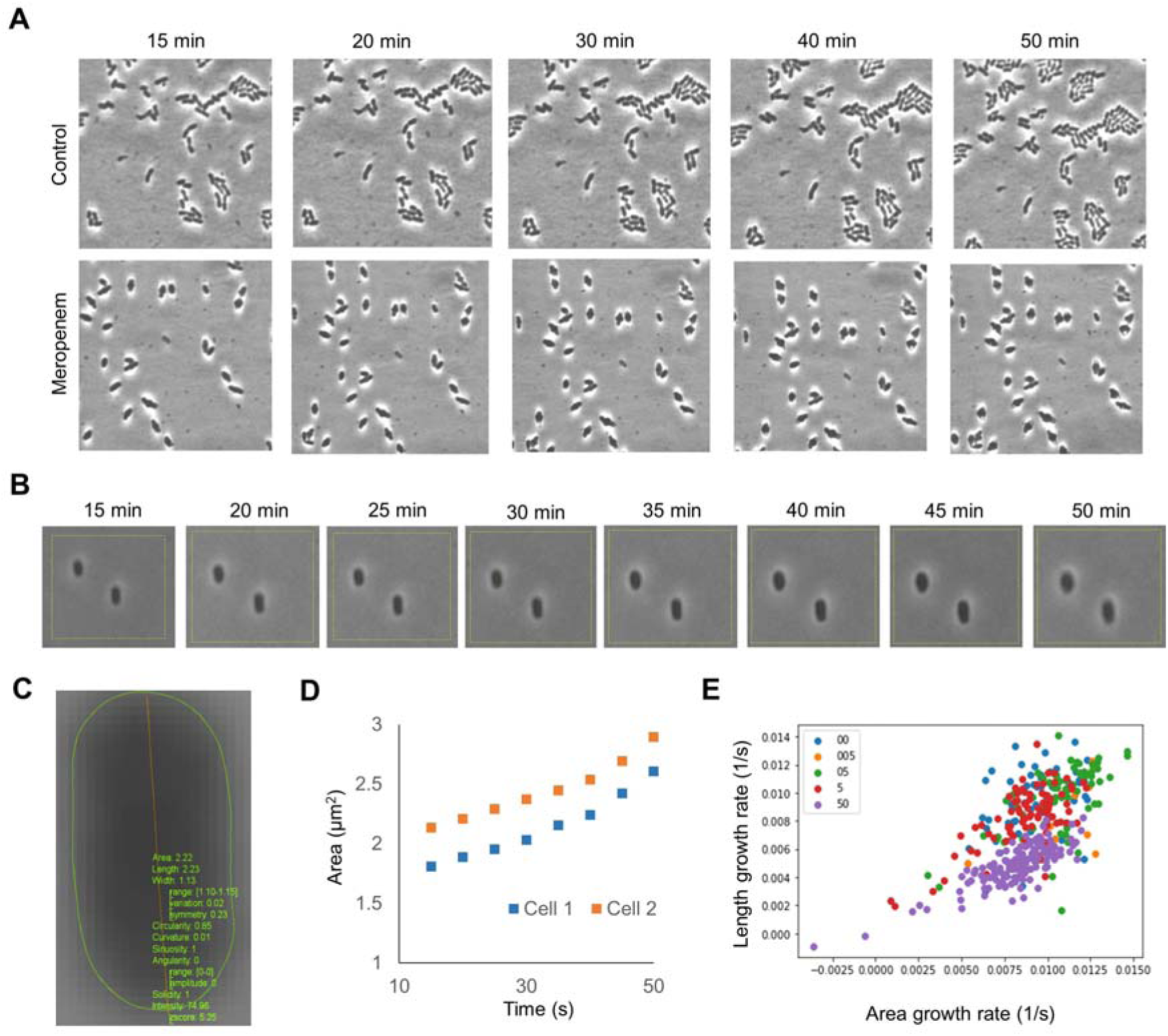
Representative results of single-cell MorphoAST of *K. pneumoniae* treated with meropenem, an intravenous β-lactam antimicrobial. (A) Time-lapse images of *K. pneumoniae* (KP0142) treated with (top) 0 and (bottom) 50 μg/ml meropenem. Time-lapse imaging was performed 15 minutes after mixing with meropenem at a 5-minute interval. The MIC of the strain was <=0.5μg/ml, and cell division was completely inhibited at 50 μg/ml. Bulging the cells was observed only in the meropenem case. (B) Zoom-in views of KP0142 treated with 50 µg/ml meropenem. (C) Morphological analysis with MicrobeJ for extracting cell features, such as area, length, width, circularity, curvature, sinuosity, angularity, and solidity. (D) Examples of two bacterial growth curves for extracting the area changing rate. (E) Distributions of area and length changing rates of individual *K. pneumoniae* exposed to various meropenem concentrations.

To automate the analysis and avoid subjectivity, the MicrobeJ plugin was applied for extracting morphological features (Figure 2C). The cell behaviors were summarized by extracting the changing rates of each feature from the data (Figure 2D and Supplementary Figure 2). Figure 2E shows an example of the distributions of length and area changing rates of a single strain (KP0142) under various meropenem concentrations. The centroids of the changing rates varied with the antimicrobial concentration. However, there were large variations among individual bacteria at the same concentrations and substantial overlaps between different concentrations. It is challenging to accurately predict the bacteria response based on one or two features. A statistical approach, specifically machine learning algorithms, is required to analyze the multiparametric data for improving classification accuracy.

### Classification of bacterial response to antimicrobial with dynamic features

We first evaluated the dynamic (or time differential) approach for predicting the bacterial response to meropenem. The dynamic features of a total of 1338 bacterial cells across various antimicrobial concentrations were measured. The cells were labeled as 1 (resistant or division) or 0 (susceptible or no division) at the strain-concentration combination based on the CDC-reported meropenem MIC for *K. pneumoniae*. Since the behaviors of individual bacteria were highly diverse, one to seven bacteria under the same strain-concentration combinations were randomly grouped. The average feature changing rates were calculated for each group. As shown in the principal component analysis (PCA) plots (Figure 3A), grouping bacteria substantially reduced the variation within a group and increased the separation between the ‘division’ and ‘no division’ groups. The separation between the groups increased with the number of bacteria in the group. The grouped data were trained and validated using the training and validation datasets with the K-Nearest Neighbors and artificial neural network classifiers (Figure 3B). For the 35-min data (i.e., 15-50 minutes of exposure), groups of one bacterium resulted in an accuracy of 89% and 90% with the K-Nearest Neighbors and artificial neural network classifiers, respectively. The prediction accuracy was generally improved by increasing the number of bacteria. With groups of seven bacteria, both classification algorithms reached over 99% accuracy. The improvement can be understood by a reduction of the statistical variation of individual cells by averaging data from multiple bacteria.

**Figure 3.**
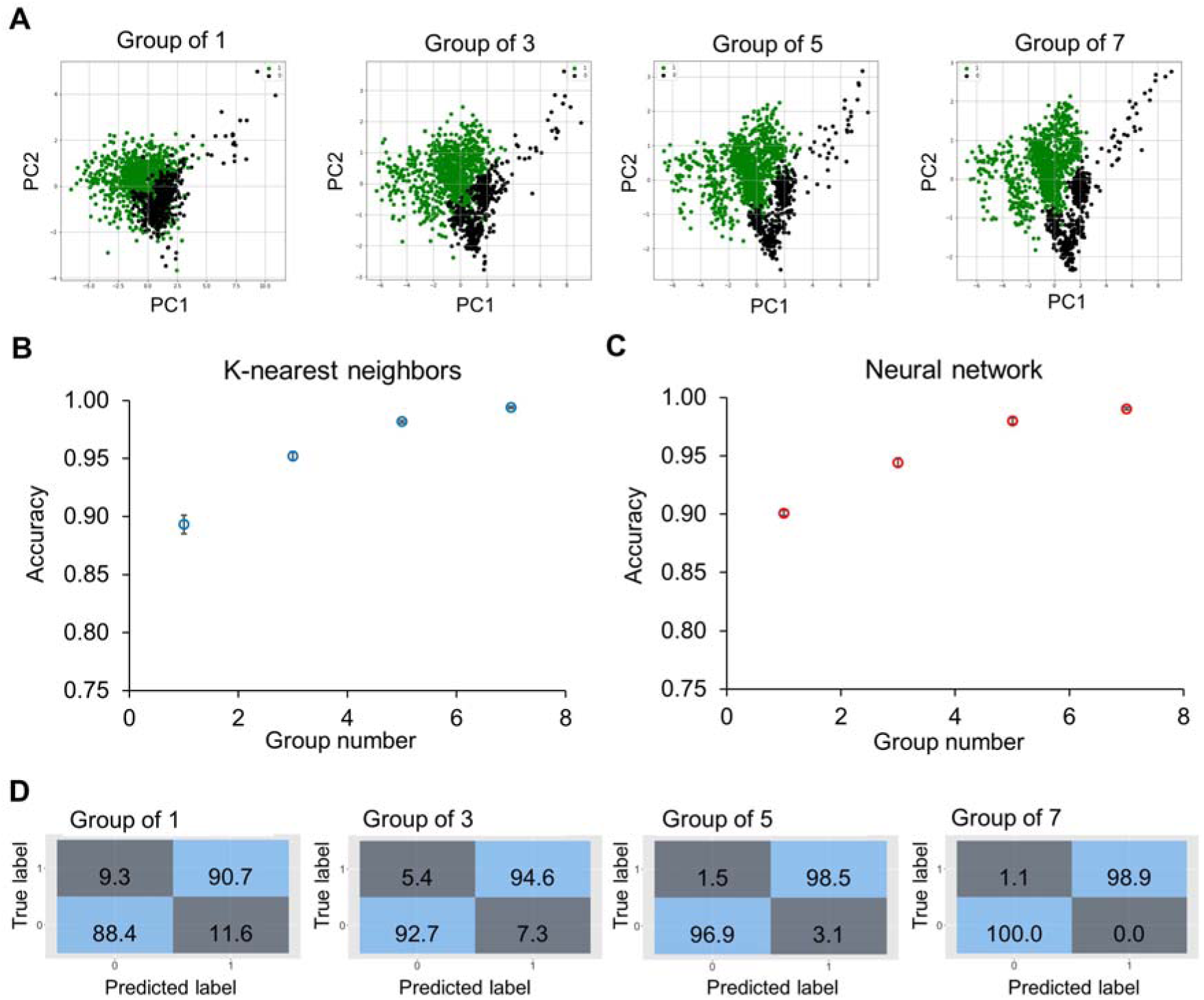
Classification accuracy of bacteria groups using the time differential approach. (A) Principal component analysis plots of dynamic features of 1338 bacteria from various bacteria strains and antimicrobial concentrations. The data are labeled as 1 (division or resistant) or 0 (no division or susceptible). Bacteria are randomly selected from the same strain-concentration combination to form a group, and the dynamic features are averaged for groups of 1, 3, 5, and 7 bacteria. (B-C) Prediction accuracy was obtained by (B) K-Nearest Neighbors and (C) artificial neural network algorithms for different group sizes. For single bacteria, the accuracy was slightly higher for the artificial neural network (∼90%). With groups of seven bacteria, both models obtained over 99% accuracy. The accuracy values are an average of 10 repetitions. (D) Evolution of confusion matrices of neural network classifiers with groups of 1 to 7 bacteria. Both the major error (true susceptible and predicted resistant) and the very major error (true resistant and predicted susceptible) were reduced with an increasing number of bacteria.

We then evaluated the accuracy of the classification model with sub-doubling times. The feature change rates were extracted using different durations of the data (5 to 35 minutes), corresponding to 15-50 minutes of antimicrobial incubation. Similarly, the data are summarized and visualized by the PCA plots (Figure 4A). The data with a 5-minute duration (i.e., between 15 minutes and 20 minutes after antimicrobial exposure) displayed a considerable variance and overlapped substantially. Nevertheless, the centroids of division and no division groups were distinct, and the separation improved with data of a longer duration. Clear separations between the groups could be observed in data with 25-min or 35-min durations. Again, we trained and validated the data using machine learning classifiers, including the K-Nearest Neighbors and artificial neural network algorithms (Figure 4B-C). These algorithms exhibited similar performances and had an accuracy of ∼80% with the 5-minute data. The results reached around 95% with the 25-minute data and over 99% with the 35-minute data. The confusion matrices indicated a false positive rate close to zero and a false negative rate of ∼1.1% (Figure 4D). These results support the use of dynamic morphological features, i.e., the time differential approach, for predicting the antimicrobial response in a sub-doubling time.

**Figure 4.**
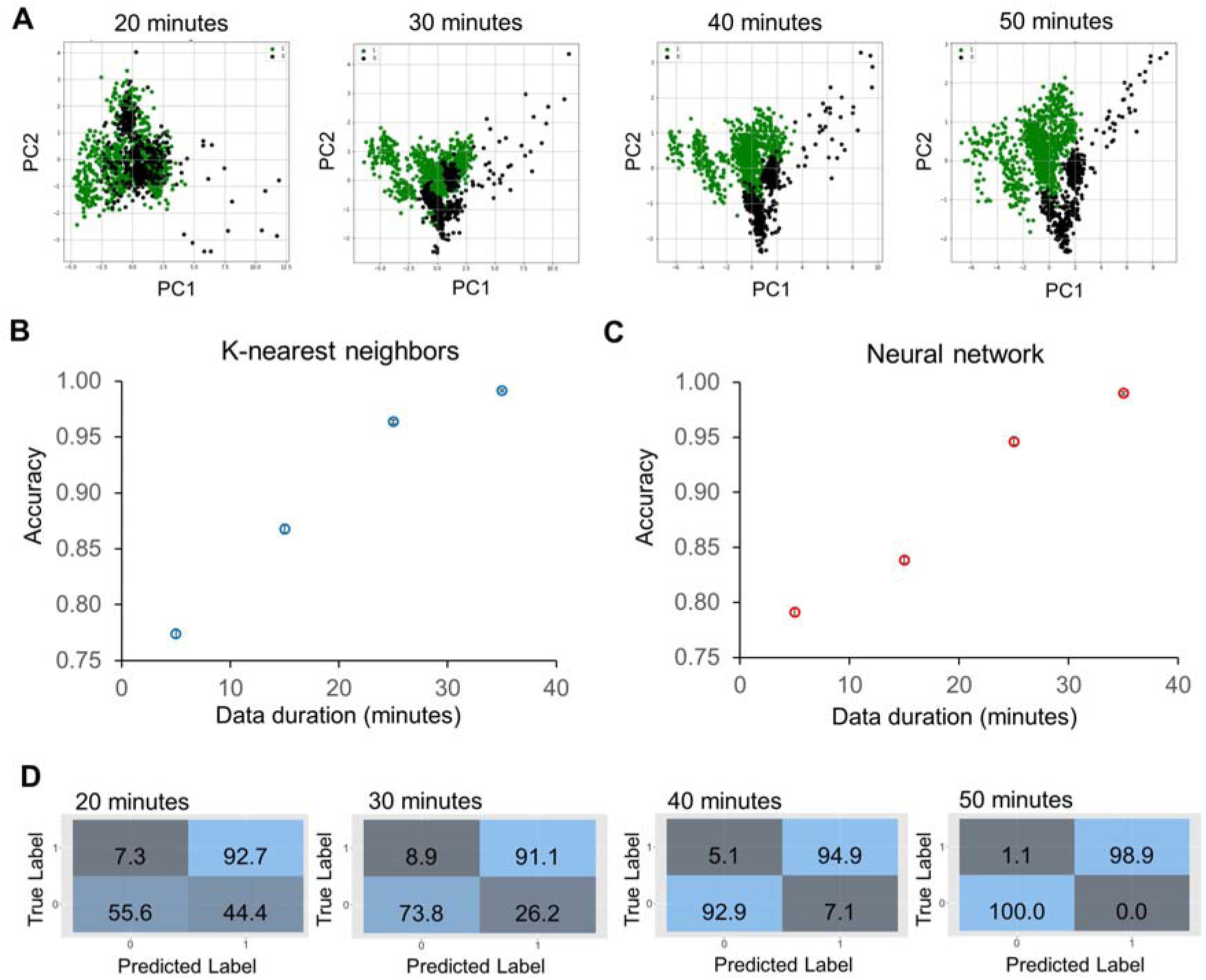
Classification accuracy of sub-doubling time susceptibility prediction using the time differential approach. (A) Principal component analysis plots of feature changing rates with 20-50 minutes of antimicrobial exposure, which correspond to 5 to 35 minutes duration of dynamic data. The data are labeled as 1 (division or resistant) or 0 (no division or susceptible) at the strain-concentration combination with groups of seven bacteria. (B-C) Prediction accuracy was obtained by (B) K-Nearest Neighbors and (C) artificial neural network algorithms for different durations of data. The accuracy was approximately 80% for a 5-min duration, and the value improved to over 99% accuracy with a 35-minute duration of data. The accuracy values are an average of 10 repetitions. (D) Evolution of confusion matrices of neural network classifiers with groups of 20-50 minutes of antimicrobial exposure. Both the major error (true susceptible and predicted resistant) and the very major error (true resistant and predicted susceptible) were reduced with the duration of the data.

### Susceptibility classification with concentration differential features

We next evaluated the concentration differential approach for predicting antimicrobial susceptibility. A total of 5050 bacterial cells with various antimicrobial concentrations were included in the prediction of survivability and susceptibility to antimicrobial exposure. The deviations of morphological features from the population means of untreated cells were calculated at each time point. Since the bacterial features against various antimicrobial concentrations were measured, the differential data were applied to classify the interpretative categories (i.e., Susceptible ‘S’, Intermediate ‘I’, and Resistant ‘R’) according to the CLSI guideline. Figure 5A shows the PCA plots of the data. The centroids of the S, I, and R groups are separated after 20 minutes of antimicrobial exposure, and there was a considerable amount of overlap. The separation widened when the antimicrobial exposure time increased. We trained and cross-validated the data with various classification algorithms, including Random Forest, Naive Bayes, K-Nearest Neighbor, and Support Vector Machine. The results suggest the Random Forest algorithm outperformed other models (Supplementary Table 3). The categorical agreement (CA), which is the percentage of interpretive agreement between the predicted and true labels, was determined (Figure 5B). The Random Forest model achieved a 97% accuracy in as early as 20 minutes (two-fifths of the doubling time) and reached over 99% accuracy in 40 minutes (four-fifths of the doubling time). Grouping multiple bacteria, however, did not improve the accuracy (Supplementary Figure 3).

**Figure 5.**
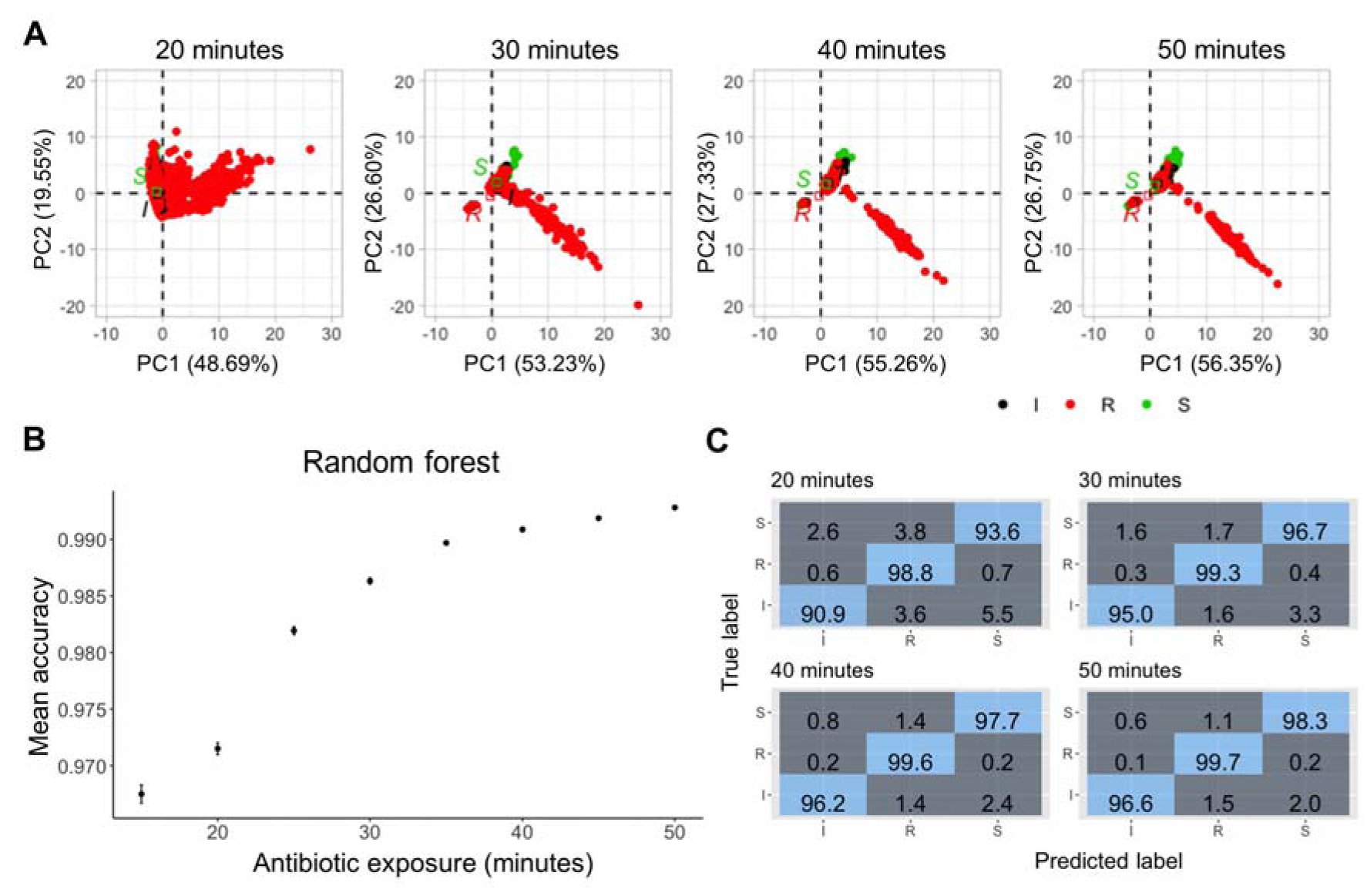
Classification accuracy of sub-doubling time susceptibility prediction using the concentration differential approach. (A) Principal component analysis plots of the normalised data with 20-50 minutes of antimicrobial exposure. The data are labeled based on the susceptibility to meropenem - red for resistant, black for intermediate, and green for susceptible. (B) Prediction accuracy of random forest classifiers with different durations of antimicrobial exposure. Median accuracy was determined using 5-fold cross-validation and 10 repetitions. Multiclass prediction accuracy was >96% as early as 15 minutes of antimicrobial exposure with increasing accuracy over time. (C) Evolution of confusion matrices of Random Forest classifiers with groups of 20-50 minutes of antimicrobial exposure. Resistant bacteria were classified with 98.8% accuracy in 20 minutes. The accuracy increased to 99.7% in 50 minutes with a very major error of 0.2% and a major error of 1.1%.

Next, the major error (ME: susceptible classified as resistant), very major error (VME: resistant classified as susceptible), and the minor errors (mE: susceptible or resistant classified as intermediate or intermediate classified as susceptible or resistant) were estimated by examining the confusion matrices (Figure 5C). With 20 minutes of meropenem exposure, the Random Forest model had a ME of 3.8%, VME of 0.7%, and a cumulative mE of 3.2%. The performance of the Random Forest model further improved with an increase in the antimicrobial exposure time. At 40 minutes of exposure, this model had a ME of 1.4%, VME of 0.2%, and a cumulative mE of 1%. These accuracy and error rates are within the FDA acceptable limits (CA>90%, VME <1.5%, and ME<3%), supporting the use of morphological features for rapid single-cell AST.

### Minimum inhibitory concentration prediction in sub-doubling times

We further evaluated the use of the concentration differential data for MIC prediction. A Random Forest regressor was trained from a total of 39,135 cells across the concentrations. The model, including historic data (i.e., earlier time points), had a cumulative improvement in performance measured by the root-mean-square error (RMSE), R-squared, and mean absolute error (MAE) values with increased exposure time (Figure 6A). The Random Forest regressor predicted the MIC in the 5-fold cross-validated training dataset with an RMSE of 0.8, MAE of 0.2, and an R^2^ of 0.93. The performance of the model was assessed against 5 unseen KP strains, which comprised 6,067 cells. The experimental MIC and predicted MIC of the 5 unseen strains based on the Random Forest regressor are compared in Figure 6B. The data showed a strong correlation, and the regressor collectively predicted the MIC within plus or minus, one two-fold dilution for all strains, resulting in an 80% essential agreement (EA) with 40 minutes of antimicrobial exposure that increases to 100% EA with 50 minutes of exposure. The predicted and experimentally reported MIC labels were also compared. Based on the predicted MIC for each strain, the model correctly predicted 100% (2/2) of the susceptible and intermediate (1/1) bacteria in as few as 30 minutes. The performance of the model increased with the antimicrobial exposure time and achieved 100% CA with 0% ME and 0% VME in 50 minutes. We then tested the model performance in predicting the MIC for individual cells within each population of the unseen strains tested (Figure 6C). While there is heterogeneity in calls for individual cells from a single population, ∼79.9% of cells from 3/5 strains tested (KP_003, KP_139, and KP_U3) had predicted MICs within one-two fold dilution from the experimental MIC after 20 minutes of antimicrobial exposure. After 50 minutes of exposure, ∼85.1% of all cells from 5/5 tested strains had predicted MICs within one-two fold dilution from experimental MIC.

**Figure 6.**
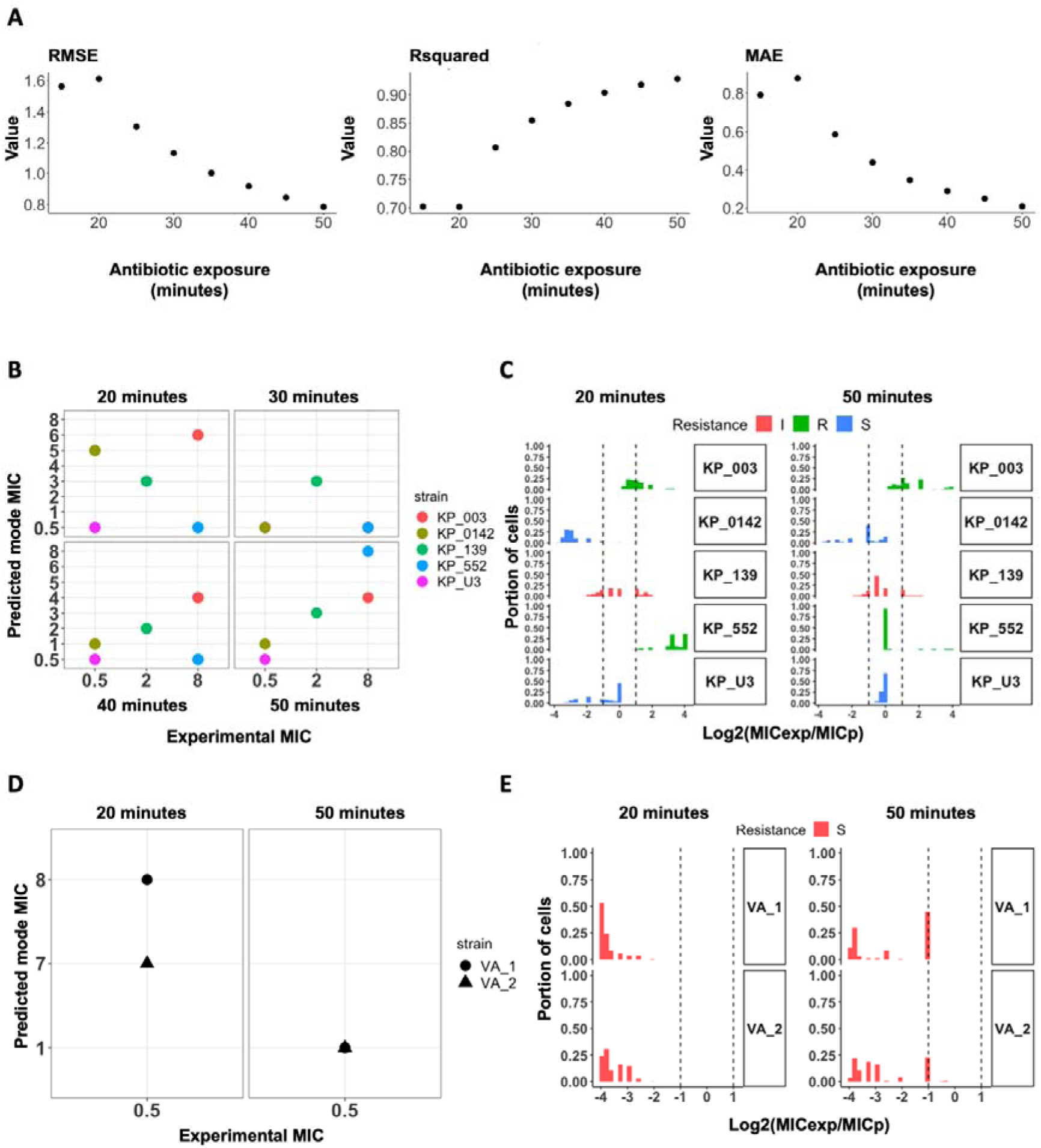
Minimum inhibitory concentration prediction with a random forest regressor using the concentration differential approach. (A) Evaluation of the regression model with root-mean-square error (RMSE), R-squared and mean absolute error (MAE) values. The regression model trained on the best set of 19 strains was assessed for the lowest RMSE and MAE and highest R-squared values. With historical data and accumulated time, the model performs better as seen in all metrics. (B) Mode of predicted MIC for each unseen test strain in comparison to the experimental MIC for each strain. The regressor achieved 100% essential agreement in the predicted mode MIC within 50 minutes. (C) Histogram of the log2 dilution of the ratio between experimental MIC (MIC_exp_) and CDC reported MIC (MIC_p_) for all the cells in the test strains. ∼85.1% of all cells had predicted MIC within one-two fold dilution from the experimental MIC after 50 minutes of exposure to meropenem. (D) MIC prediction and susceptibility classification of two clinical isolates from patients with urinary tract infections with 922 (VA_1) and 648 (VA_2) cells imaged across multiple concentrations of meropenem. The model predicts the mode MIC for VA_1 and VA_2 to be 1 µg/mL and accurately classified both to be susceptible. (E) Histogram of the log2 dilution of the ratio between reported MIC (MIC_rep_) and CDC reported MIC (MIC_pred_). Extensive heterogeneity in the populations of the clinical isolates, even within 50 minutes, only ∼40% of the cells from the two isolates starts to show susceptibility phenotype.

In addition to validation against unseen strains, the model was tested against imaging data from two clinical samples of KP obtained from patients visiting the VA (Palo Alto) with urinary tract infections (Figure 6D-E). The experimental MICs for these clinical samples (VA_1 and VA_2) was 0.5 µg/mL, i.e., susceptible. Within 20 minutes of meropenem treatment, the model did not observe patterns for susceptibility it had learned from the training data and predicted both strains to be resistant with an MIC of 8 and 7 µg/mL respectively. When the incubation time was increased to 50 minutes, both strains were predicted to have a mode MIC of 1 µg/mL which accurately classifies both clinical isolates as susceptible with an MIC within one-two fold dilution from the experimental MIC resulting. Notably, heterogeneity within the population is more readily apparent with the clinical isolates where even after the 50 minutes of exposure, only a subset of the population started portraying susceptibility features as determined by the predicted MIC - ∼45% of VA_1 and ∼23.3% of VA_2 had predicted MICs within one-two fold dilution with the mode MIC of 1 µg/mL. These results suggest a sufficient number of bacteria should be considered to compensate the heterogeneity of clinical samples in order to accurately predict the MIC.^30^

## Discussion

This study demonstrated a rapid workflow for determining the antimicrobial susceptibility of *K. pneumoniae* against meropenem using time-lapse single-cell imaging, computer vision, and machine learning models. By tracking the antimicrobial-induced morphological changes of individual cells, the MorphoAST workflow predicted the susceptibility category and MIC with high accuracy in a fraction of the doubling time of the bacteria. We evaluated two data processing strategies based on the time and concentration differentials. Both strategies successfully predicted the susceptibility in as few as 20 minutes (two-fifths of the doubling time) of antimicrobial exposure with high accuracy. The concentration differential approach with the random forest classifier achieved an overall better accuracy and resolution for predicting the susceptibility category and MIC (CA >97% in as few as 20 minutes). Therefore, the concentration differential approach should be applied whenever possible. The time differential approach, however, could be useful when only a small number of bacteria is available (e.g., direct detection of bacteria from clinical samples) and only limited antimicrobial concentrations can be measured (e.g., a point-of-care device that detects only a small number of conditions). Our data also suggested that the prediction accuracy was generally improved with the antimicrobial exposure time, the number of bacteria being analyzed, and the number of testing conditions. These results underscore important considerations and tradeoffs in the design of the single-cell AST workflow and provide examples of future experimental designs.^31^

Our results showed that morphological features are useful information for rapid AST. In particular, the formation of bulges among *K. pneumoniae* with varying meropenem MIC are distinct and allowed us to classify susceptibility in a sub-doubling time. Bulge formation has been associated with the disruption of peptidoglycan and cell wall degradation before cell lysis.^24^) Prior studies have also reported the formation of cell-wall deficient spheroplasts in carbapenem-tolerant *K. pneumoniae* strains, after exposure to meropenem.^32,33^ While antimicrobial-induced shape changes in bacterial cells are not fully understood in general, recent quantitative modeling reveals potential advantages of this physiological adaptation, which include decreasing antimicrobial influx and diluting intracellular antimicrobials, leading to a higher tolerance.^34^ The size and shape regulation differs from one organism to another, but importantly, provides a general reflection of their response in a short time frame.^35^ As demonstrated in this study, MorphoAST provides a workflow for utilizing the potential of morphological features for rapid AST. The workflow should be compatible with various morphological features, such as filamentation, bulging, and lysis, induced by specific antimicrobial-bacteria combinations. The methodical procedure identifies and distinguishes key morphological features that can be captured before and after the treatment (time differential) or with various antimicrobial dilutions (concentration differential).

A major advantage of the MorphoAST workflow is the short turnaround time. As the morphological features can be captured in a fraction of the doubling time, it dramatically reduces the assay time compared to the standard phenotypic AST (e.g., broth microdilution), which typically requires one or more days. The approach bypasses the requirement of cell replication in other single-cell AST techniques.^36^ This characteristic will be particularly useful for diagnosing slow-growing and difficult-to-culture bacteria in normal laboratory conditions. Compared to other single-cell analysis techniques that measure the nanoscale motion of bacteria and metabolic activities,^26–28^ the MorphoAST workflow requires only a small inoculum size and a relatively simple setup consisting mainly of agar pads and a microscope. As imaging is common in AST^37,38^ and low-cost microscopes are readily available, the MorphoAST workflow can be integrated with existing systems and implemented in a variety of settings. These advantages and characteristics will potentially increase the utility of MorphoAST for direct sample AST testing, especially in critical diseases like sepsis where the bacteria load in the blood is typically very low. The small inoculum will also considerably reduce the time to AST results at the point of care.

The extension of MorphoAST upon future validation with other drug-bug combinations will potentially accelerate microbiological analysis for combating multidrug-resistant bacteria. In both unseen cells and clinical isolates, our results demonstrate outstanding categorical agreement (100%) and essential agreement (100%), which are above the FDA acceptable level (≥90%) for new AST devices. The machine learning models consistently had 0% VME after 50 minutes of antimicrobial exposure, which is ideal and substantially better than the acceptable VME rate (≤1.5%). A low VME rate is particularly important as missing a resistant strain could result in the prescription of an ineffective antimicrobial for the patient. More investigation will be required to elucidate the mechanisms of action of meropenem and its effects on other bacteria. Further optimization by incorporating population dynamics into the training methodology could shorten the time to decision and potentially increase the accuracy with smaller population sizes. Bacterial populations are heterogenous, and cells do not all portray susceptibility phenotypes simultaneously, and hence, deciphering the dynamics of a population trending towards susceptibility early can accelerate AST further. Towards clinical translation of the proof-of-concept MorphoAST, additional drug-bug combinations should be tested to evaluate the accuracy of the approach in other settings. Automation of the drug mixing and bacteria trapping steps will shorten the initial preparation time and capture changes in bacterial morphologies at earlier time points. Implementing the workflow in an integrated, low-cost imaging system, instead of a microscope, will also be important for disseminating the workflow for managing a wide spectrum of infectious diseases.

## Methods

### Bacteria culture

*Klebsiella pneumoniae* isolates (Supplementary Table 1) from the CDC & FDA Antimicrobial Resistance (AR) bank, Johns Hopkins University, and American Type Culture Collective (ATCC) were grown in Mueller Hinton Broth (MHB) at 37°C overnight. The next day, cells were sub-cultured in fresh MHB media for three hours (growth phase) to a density equivalent to OD_600_ = 0.5. Bacteria cells (10 μL) were treated with 10 μL meropenem at varying final concentrations of 0 (control), 0.05, 0.5, 5, and 50 µg/mL to ascertain the physiological responses of cells. A subset of these isolates was treated with finer range of antimicrobial concentrations (0.02, 0.04, 0.016, 0.032, 0.064, 0.125, 0.5, 1, 2 µg/mL) to train the algorithm for quantitative prediction of MIC. The MICs of the isolates were experimentally obtained and confirmed in triplicate using broth microdilution method according to CSLI guidelines.

### Bacteria imaging

Bacteria cells and meropenem in liquid media are briefly vortexed before mounting on a 1% (v/v) UltraPure agarose pad (10 x 20 mm, Invitrogen US) in M9 minimal media (Gibco, US) on a micro slide covered with a glass coverslip (#1.5, 22 x 30 mm). Bacteria imaging was performed using a Nikon Ti2-E inverted microscope equipped with a DS-Qi2 CMOS camera and an Okolab stage-top temperature control chamber (37°C, 5% CO_2_). Images were acquired using a Nikon CFI Plan Apochromat l DM 100X oil objective lens and an external Phase Contrast (Ph3) module. Each isolate was observed over a period of 75 minutes (15-90 minutes after mixing with the antimicrobial).

### Feature extraction

Time-series images were stacked and corrected for shifts in the time series using the Template Matching and Slice Alignment ImageJ plugin.^39^ Each stack was then analyzed using the MicrobeJ plugin for ImageJ.^29^ The plugin, designed for the detection and analysis of bacterial cells, uses computer vision algorithms to automatically identify a cell and determine a suite of morphological features, such as its area, length, circularity, and perimeter (Supplementary Table 2). The tracking data were summarized as a table and exported as a .csv file for further analysis.

### Data analysis and machine learning

The morphological data were analyzed to predict the susceptibility and MIC of the bacterial strain before the average doubling time (∼50 minutes). Two differential strategies based on the time-dependent or antimicrobial concentration-dependent changes of the bacteria were applied. The processed data were then applied for susceptibility classification and MIC prediction. A schematic of the workflow is shown in Figure 1.

In the time differential (or dynamic) approach, processed image stacks for *Klebsiella pneumoniae* isolates (KP016, KP0140, KP1705, KP0153, and KP0142) at various antimicrobial concentrations were evaluated. The cells were labeled as division (resistant) or no division (susceptible). Data analyses were performed using Python with data analysis and machine learning libraries.^40,41^ To explore AST in a sub-doubling time, time-lapse data from the first 35 minutes (corresponding to 15 to 50 minutes after antimicrobial exposure) at a 5-minute interval were analyzed. Erroneous data (e.g., missing bacteria) were detected and removed. Feature-changing rates of individual cells were determined by fitting the time-lapse data with the exponential function. The area, aspect ratio, circularity, length, and perimeter were the most relevant dynamic features. The changing rates of these features for each cell, along with the meropenem concentration, time of tracking, the bacterial strain, and the label for growing/not growing, were used to train the K-Nearest Neighbor and artificial neural network (MPLClassifier) models from the scikit-learn machine learning library.^42^ For the K-Nearest Neighbor model, k was optimal at 20. The optimal structure of the artificial neural network was found at three hidden layers with five neurons each, even though a simple grid (two hidden layers with two neurons each) was sufficient for well-separated data (e.g., 50-minute data). The accuracy of each model was obtained by averaging the values of 10 runs.

In the concentration differential approach, the effect of each concentration of meropenem on individual cells was determined as a deviation from the population mean of untreated cells at the same time point for each of the 19 features, concentrations, and strains. Only cells detected within the first three time points and with no missing data were included in the analysis. If a cell died within the observation timeframe, the subsequent time points were marked with zero values for the features. The best modeling strategy for the classification of the normalized data was chosen among Random Forest, Naive Bayes, K-Nearest Neighbor, and Support Vector Machine based on the accuracy of models trained at the first time point (Supplementary Table 3). Random Forest classification models were trained on the normalized dataset for each time point with 5-fold cross-validation and 10 repetitions using the R package caret.^43^) The models with the highest accuracy were retained.

To develop the regression model for MIC prediction, processed image stacks for 24 *K. pneumoniae* strains that spanned 10 resistant, 12 susceptible, and 2 intermediate against meropenem were used. We searched for an optimal training set of 19 strains that encompasses the diverse morphological changes and predicts the MIC and susceptibility of the other unseen 2 resistant, 2 susceptible, and 1 intermediate KP strains through 30 iterations of the possible combinations. The data were normalized as described above in the random forest. The best modeling strategy for the prediction of MIC was chosen among Linear Regression, Neural Net, Random Forest, K-Nearest Neighbor, Support Vector Machine, and Gaussian Process Regression based on the lowest root mean square error (RMSE) and highest coefficient of determination (R^2^) for models trained at the first time point (Supplementary Table 4). The training data were weighted to account for any class-level imbalance by dividing the number of cells in the intermediate category by the number of cells in the corresponding susceptible, resistant, and intermediate categories. Random Forest regression models were trained for each time point with 5-fold cross-validation and 5 repetitions using R packages ranger^44^ and caret. ^43^ Models were trained against the experimental MIC for these strains, and the model with the lowest RMSE, mean absolute error (MAE), and highest R^2^ was selected. The final models were tested against the 5 left out unseen strains and an additional 2 clinical isolates from patients with UTI. Mode MIC for the population of cells for each strain was attributed as the MIC for the strain. Categorical agreement (CA) and essential agreement (EA) for the cross-validated test as well as the left-out test were determined. The major error (ME) and very major error (VME) rates were calculated for the model. The model with the best performance against unseen strains was selected.

### Clinical urine samples

Two de-identified, remaindered urine samples (10 mL) from patients infected with *Klebsiella pneumoniae* were collected from the Veterans Affairs Palo Alto Health Care System clinical microbiology laboratory. Klebsiella was identified by the clinical lab and confirmed by plating on ChromAgar Orientation (BD). The urine samples were stored in the −80 °C freezer until used. Next, the urine samples were streaked on *ChromoSelect* selective agar (Millipore) and grown overnight to obtain isolated colonies. The MIC for these strains against meropenem was determined through micro-broth dilution in triplicate. The isolates were grown, treated with meropenem, and imaged as described above.

### Role of funders

The funders of this study were not involved in the study design, data collection, analysis, interpretation and writing of the report.

## Contributors

Conceptualization: S.Y., P.K.W., K.C.T., N.R., M.R. which was refined by input from J.C.L and K.E.M. Methodology: S.Y., P.K.W., K.C.T., N.R., M.R. Investigation: K.C.T., N.R., M.R., E.Z. Imaging: K.C.T., E.Z. Data analysis: K.C.T., N.R., M.R., Z.Q., O.E. Funding acquisition: S.Y., P.K.W., J.C.L. Project administration: S.Y., P.K.W., J.C.L. Supervision: S.Y., P.K.W., J.C.L. Writing – original draft: P.K.W., K.C.T., N.R., M.R. All authors discussed the results and contributed to reviewing and editing the manuscript.

## Declaration of interests

Authors declare that they have no competing interests.

## Acknowledgment

We acknowledge technical support from Youngbin Lim from the Stanford Cell Sciences Imaging Facility. We are grateful for the insightful discussions and computational support from Dean Deng from Stanford University. This work was performed, in part, at the Stanford Cell Sciences Imaging Facility of Stanford University. All computational analyses were performed on the Stanford Genomic Cluster of the Stanford Research Computing Center.

## Funding

This work was supported in part by NIH NIAID (R01AI153133).

## Data sharing and materials availability

The data, code, and materials that support the findings of this study will be openly available in a public repository and from the corresponding authors PW and SY on request.

## Notes

### Competing Interest Statement

The authors have declared no competing interest.

### Summary of Updates

inclusion of clinical samples, additional bacterial strains, as well as an optimized training dataset

## References

1. Nordmann, P.; Naas, T.; Poirel, L. Global Spread of Carbapenemase Producing Enterobacteriaceae. Emerg Infect Dis 2011, 17 (10), 1791–1798. https://doi.org/10.3201/eid1710.110655.

2. Durante-Mangoni, E.; Andini, R.; Zampino, R. Management of Carbapenem-Resistant Enterobacteriaceae Infections. Clinical Microbiology and Infection 2019, 25 (8), 943–950. https://doi.org/10.1016/j.cmi.2019.04.013.

3. Jernigan, J. A.; Hatfield, K. M.; Wolford, H.; Nelson, R. E.; Olubajo, B.; Reddy, S. C.; McCarthy, N.; Paul, P.; McDonald, L. C.; Kallen, A.; Fiore, A.; Craig, M.; Baggs, J. Multidrug-Resistant Bacterial Infections in U.S. Hospitalized Patients, 2012–2017. New England Journal of Medicine 2020, 382 (14), 1309–1319. https://doi.org/10.1056/nejmoa1914433.

4. Jorgensen, J. H.; Ferraro, M. J. Antimicrobial Susceptibility Testing: A Review of General Principles and Contemporary Practices. Clinical Infectious Diseases 2009, 49 (11), 1749–1755. https://doi.org/10.1086/647952.

5. Vasala, A.; Hytönen, V. P.; Laitinen, O. H. Modern Tools for Rapid Diagnostics of Antimicrobial Resistance. Front Cell Infect Microbiol 2020, 10, 308. https://doi.org/10.3389/fcimb.2020.00308.

6. Tjandra, K. C.; Ram-mohan, N.; Abe, R.; Hashemi, M. M.; Lee, J.; Chin, S. M.; Roshardt, M. A.; Liao, J. C.; Wong, P. K.; Yang, S. Diagnosis of Bloodstream InfectionsLJ: An Evolution of Technologies towards Accurate and Rapid Identification and Antibiotic Susceptibility Testing. 2022.

7. Baltekin, Ö.; Boucharin, A.; Tano, E.; Andersson, D. I.; Elf, J. Antibiotic Susceptibility Testing in Less than 30 Min Using Direct Single-Cell Imaging. Proc Natl Acad Sci U S A 2017, 114 (34), 9170–9175. https://doi.org/10.1073/pnas.1708558114.

8. Li, H.; Torab, P.; Mach, K. E.; Surrette, C.; England, M. R.; Craft, D. W.; Thomas, N. J.; Liao, J. C.; Puleo, C.; Wong, P. K. Adaptable Microfluidic System for Single-Cell Pathogen Classification and Antimicrobial Susceptibility Testing. Proceedings of the National Academy of Sciences 2019, 201819569. https://doi.org/10.1073/pnas.1819569116.

9. Choi, J.; Yoo, J.; Lee, M.; Kim, E. G.; Lee, J. S.; Lee, S.; Joo, S.; Song, S. H.; Kim, E. C.; Lee, J. C.; Kim, H. C.; Jung, Y. G.; Kwon, S. A Rapid Antimicrobial Susceptibility Test Based on Single-Cell Morphological Analysis. Sci Transl Med 2014, 6 (267). https://doi.org/10.1126/scitranslmed.3009650.

10. Lu, Y.; Gao, J.; Zhang, D. D.; Gau, V.; Liao, J. C.; Wong, P. K. Single Cell Antimicrobial Susceptibility Testing by Confined Microchannels and Electrokinetic Loading. Anal Chem 2013, 85 (8), 3971–3976. https://doi.org/10.1021/ac4004248.

11. Zhang, F.; Jiang, J.; McBride, M.; Yang, Y.; Mo, M.; Iriya, R.; Peterman, J.; Jing, W.; Grys, T.; Haydel, S. E.; Tao, N.; Wang, S. Direct Antimicrobial Susceptibility Testing on Clinical Urine Samples by Optical Tracking of Single Cell Division Events. Small 2020, 16 (52), 1–10. https://doi.org/10.1002/smll.202004148.

12. Choi, J.; Jung, Y. G.; Kim, J.; Kim, S.; Jung, Y.; Na, H.; Kwon, S. Rapid Antibiotic Susceptibility Testing by Tracking Single Cell Growth in a Microfluidic Agarose Channel System. Lab Chip 2013, 13 (2), 280–287. https://doi.org/10.1039/c2lc41055a.

13. Łapińska, U.; Voliotis, M.; Lee, K. K.; Campey, A.; Stone, M. R. L.; Tuck, B.; Phetsang, W.; Zhang, B.; Tsaneva-Atanasova, K.; Blaskovich, M. A.; Pagliara, S. Fast Bacterial Growth Reduces Antibiotic Accumulation and Efficacy. Elife 2022, 11, 1–33. https://doi.org/10.7554/elife.74062.

14. Brauner, A.; Fridman, O.; Gefen, O.; Balaban, N. Q. Distinguishing between Resistance, Tolerance and Persistence to Antibiotic Treatment. Nat Rev Microbiol 2016, 14 (5), 320–330. https://doi.org/10.1038/nrmicro.2016.34.

15. Veses-Garcia, M.; Antypas, H.; Löffler, S.; Brauner, A.; Andersson-Svahn, H.; Richter-Dahlfors, A. Rapid Phenotypic Antibiotic Susceptibility Testing of Uropathogens Using Optical Signal Analysis on the Nanowell Slide. Front Microbiol 2018, 9 (JUL), 1–10. https://doi.org/10.3389/fmicb.2018.01530.

16. Matsumoto, Y.; Sakakihara, S.; Grushnikov, A.; Kikuchi, K.; Noji, H.; Yamaguchi, A.; Iino, R.; Yagi, Y.; Nishino, K. A Microfluidic Channel Method for Rapid Drug-Susceptibility Testing of Pseudomonas Aeruginosa. PLoS One 2016, 11 (2), 1–17. https://doi.org/10.1371/journal.pone.0148797.

17. Avesar, J.; Rosenfeld, D.; Truman-Rosentsvit, M.; Ben-Arye, T.; Geffen, Y.; Bercovici, M.; Levenberg, S. Rapid Phenotypic Antimicrobial Susceptibility Testing Using Nanoliter Arrays. Proc Natl Acad Sci U S A 2017, 114 (29), E5787–E5795. https://doi.org/10.1073/pnas.1703736114.

18. Kalashnikov, M.; Mueller, M.; McBeth, C.; Lee, J. C.; Campbell, J.; Sharon, A.; Sauer-Budge, A. F. Rapid Phenotypic Stress-Based Microfluidic Antibiotic Susceptibility Testing of Gram-Negative Clinical Isolates. Sci Rep 2017, 7 (1), 1–10. https://doi.org/10.1038/s41598-017-07584-z.

19. Bourne, C. R. Bacterial Growth Mindset: Structural Plasticity in Defense Systems. Structure 2021, 29 (2), 97–98. https://doi.org/10.1016/j.str.2021.01.007.

20. Yang, D. C.; Blair, K. M.; Salama, N. R. Staying in Shape: The Impact of Cell Shape on Bacterial Survival in Diverse Environments. Microbiology and Molecular Biology Reviews 2016, 80 (1), 187–203. https://doi.org/10.1128/mmbr.00031-15.

21. Bruus, H. Acoustofluidics 10: Scaling Laws in Acoustophoresis. Lab Chip 2012, 12 (9), 1578. https://doi.org/10.1039/c2lc21261g.

22. Zahir, T.; Camacho, R.; Vitale, R.; Ruckebusch, C.; Hofkens, J.; Fauvart, M.; Michiels, J. High-Throughput Time-Resolved Morphology Screening in Bacteria Reveals Phenotypic Responses to Antibiotics. Commun Biol 2019, 2 (1), 1–13. https://doi.org/10.1038/s42003-019-0480-9.

23. Nonejuie, P.; Burkart, M.; Pogliano, K.; Pogliano, J. Bacterial Cytological Profiling Rapidly Identifies the Cellular Pathways Targeted by Antibacterial Molecules. Proc Natl Acad Sci U S A 2013, 110 (40), 16169–16174. https://doi.org/10.1073/pnas.1311066110.

24. Yao, Z.; Kahne, D.; Kishony, R. Distinct Single-Cell Morphological Dynamics under Beta-Lactam Antibiotics. Mol Cell 2012, 48 (5), 705–712. https://doi.org/10.1016/j.molcel.2012.09.016.

25. Monahan, L. G.; Turnbull, L.; Osvath, S. R.; Birch, D.; Charles, I. G.; Whitchurch, C. B. Rapid Conversion of Pseudomonas Aeruginosa to a Spherical Cell Morphotype Facilitates Tolerance to Carbapenems and Penicillins but Increases Susceptibility to Antimicrobial Peptides. Antimicrob Agents Chemother 2014, 58 (4), 1956–1962. https://doi.org/10.1128/AAC.01901-13.

26. Rosłoń, I. E.; Japaridze, A.; Steeneken, P. G.; Dekker, C.; Alijani, F. Probing Nanomotion of Single Bacteria with Graphene Drums. Nat Nanotechnol 2022, 17 (6), 637–642. https://doi.org/10.1038/s41565-022-01111-6.

27. Scherer, B.; Surrette, C.; Li, H.; Torab, P.; Kvam, E.; Galligan, C.; Go, S.; Grossmann, G.; Hammond, T.; Johnson, T.; St-Pierre, R.; Nelson, J. R.; Potyrailo, R. A.; Khire, T.; Hsieh, K.; Wang, T. H.; Wong, P. K.; Puleo, C. M. Digital Electrical Impedance Analysis for Single Bacterium Sensing and Antimicrobial Susceptibility Testing. Lab Chip 2021, 21 (6), 1073–1083. https://doi.org/10.1039/d0lc00937g.

28. Kaushik, A. M.; Hsieh, K.; Mach, K. E.; Lewis, S.; Puleo, C. M.; Carroll, K. C.; Liao, J. C.; Wang, T. H. Droplet-Based Single-Cell Measurements of 16S RRNA Enable Integrated Bacteria Identification and Pheno-Molecular Antimicrobial Susceptibility Testing from Clinical Samples in 30 Min. Advanced Science 2021, 8 (6), 1–14. https://doi.org/10.1002/advs.202003419.

29. Ducret, A.; Quardokus, E. M.; Brun, Y. V. MicrobeJ, a Tool for High Throughput Bacterial Cell Detection and Quantitative Analysis. Nat Microbiol 2016, 1 (7), 1–7. https://doi.org/10.1038/nmicrobiol.2016.77.

30. Smith, K. P.; Kirby, J. E. The Inoculum Effect in the Era of Multidrug Resistance: Minor Differences in Inoculum Have Dramatic Effect on MIC Determination. Antimicrob Agents Chemother 2018, 62 (8). https://doi.org/10.1128/AAC.00433-18.

31. Li, H.; Hsieh, K.; Wong, P. K.; Mach, K. E.; Liao, J. C.; Wang, T. H. Single-Cell Pathogen Diagnostics for Combating Antibiotic Resistance. Nature Reviews Methods Primers 2023, 3 (1). https://doi.org/10.1038/s43586-022-00190-y.

32. Cross, T.; Ransegnola, B.; Shin, J.-H.; Weaver, A.; Fauntleroy, K.; VanNieuwenhze, M. S.; Westblade, L. F.; Dörr, T. Spheroplast-Mediated Carbapenem Tolerance in Gram-Negative Pathogens. Antimicrob Agents Chemother 2019, 63 (9). https://doi.org/10.1128/AAC.00756-19.

33. Murtha, A. N.; Kazi, M. I.; Schargel, R. D.; Cross, T.; Fihn, C.; Cattoir, V.; Carlson, E. E.; Boll, J. M.; Dörr, T. High-Level Carbapenem Tolerance Requires Antibiotic-Induced Outer Membrane Modifications. PLoS Pathog 2022, 18 (2), 1–19. https://doi.org/10.1371/journal.ppat.1010307.

34. Ojkic, N.; Serbanescu, D.; Banerjee, S. Antibiotic Resistance via Bacterial Cell Shape-Shifting. mBio 2022. https://doi.org/10.1128/mbio.00659-22.

35. Ojkic, N.; Banerjee, S. Bacterial Cell Shape Control by Nutrient-Dependent Synthesis of Cell Division Inhibitors. Biophys J 2021, 120 (11), 2079–2084. https://doi.org/10.1016/j.bpj.2021.04.001.

36. Pancholi, P.; Carroll, K. C.; Buchan, B. W.; Chan, R. C.; Dhiman, N.; Ford, B.; Granato, P. A.; Harrington, A. T.; Hernandez, D. R.; Humphries, R. M.; Jindra, M. R.; Ledeboer, N. A.; Miller, S. A.; Brian Mochon, A.; Morgan, M. A.; Patel, R.; Schreckenberger, P. C.; Stamper, P. D.; Simner, P. J.; Tucci, N. E.; Zimmerman, C.; Wolk, D. M. Multicenter Evaluation of the Accelerate PhenoTest BC Kit for Rapid Identification and Phenotypic Antimicrobial Susceptibility Testing Using Morphokinetic Cellular Analysis. J Clin Microbiol 2018, 56 (4). https://doi.org/10.1128/JCM.01329-17.

37. Song, D.; Lei, Y. Mini-Review: Recent Advances in Imaging-Based Rapid Antibiotic Susceptibility Testing. Sensors and Actuators Reports 2021, 3 (October). https://doi.org/10.1016/j.snr.2021.100053.

38. Salido, J.; Bueno, G.; Ruiz-santaquiteria, J.; Cristobal, G. A Review on Low-Cost Microscopes for Open Science. 2022, No. June, 1–14. https://doi.org/10.1002/jemt.24200.

39. Qingzong, T. Template Matching and Slice Alignment. https://sites.google.com/site/qingzongtseng/template-matching-ij-plugin#credit (accessed 2022-06-28).

40. Wes McKinney. Python for Data Analysis; 2017; Vol. 71. https://doi.org/10.1097/00007890-200105270-00005.

41. Harris, C. R.; Millman, K. J.; van der Walt, S. J.; Gommers, R.; Virtanen, P.; Cournapeau, D.; Wieser, E.; Taylor, J.; Berg, S.; Smith, N. J.; Kern, R.; Picus, M.; Hoyer, S.; van Kerkwijk, M. H.; Brett, M.; Haldane, A.; del Río, J. F.; Wiebe, M.; Peterson, P.; Gérard-Marchant, P.; Sheppard, K.; Reddy, T.; Weckesser, W.; Abbasi, H.; Gohlke, C.; Oliphant, T. E. Array Programming with NumPy. Nature 2020, 585 (7825), 357–362. https://doi.org/10.1038/s41586-020-2649-2.

42. Pedregosa, F.; Michel, V.; Grisel, O.; Blondel, M.; Prettenhofer, P.; Weiss, R.; Vanderplas, J.; Cournapeau, D.; Varoquaux, G.; Gramfort, A.; Thirion, B.; Grisel, O.; Dubourg, V.; Passos, A.; Brucher, M.; Perrot, M.; Duchesnay, É. Scikit-Learn: Machine Learning in Python. Journal of Machine Learning Research 2011, 12, 2825–2830. https://doi.org/10.5555/1953048.2078195.

43. Kuhn, M. Building Predictive Models in R Using the Caret Package. J Stat Softw 2008, 28 (5), 1–26. https://doi.org/10.18637/jss.v028.i05.

44. Wright, M. N.; Ziegler, A. RangerLJ: A Fast Implementation of Random Forests for High Dimensional Data in C++ and R. J Stat Softw 2017, 77 (1). https://doi.org/10.18637/jss.v077.i01.

