## Supplemental Information for "Machine Learning Workflow for Single-Cell Antimicrobial Susceptibility Testing of *Klebsiella pneumoniae* to Meropenem in Sub-Doubling Time"

[#] These authors contributed equally

**Supplementary Table 1.** Klebsiella pneumoniae strains are clinical isolates from CDC/FDA Antimicrobial Resistance Isolate Bank, ATCC, Johns Hopkins University, and Palo Alto Veterans Affairs Hospital. Experimental MICs were verified in triplicate utilizing microbroth dilution.

| No | KP Strains | Meropenem Susc. | Experimental MIC | Reported MIC | Reported Resistance Mechanisms | Sequence Accession # |
|---|---|---|---|---|---|---|
| **CDC Antibiotic Resistance Isolate Bank** | | | | | | |
| 1 | AR_0003 | R | >8 | >8 | aac(6')-Ib, aadA1, aph(3')-Ib, aph(6)-Id, dfrA14, EMRD, KDEA, **KPC-3**, Omp35, OmpK35, oqxA, oqxB, OXA-9, SHV-12, sul2, TEM-1A | SAMN04014844 |
| 2 | AR_0004 | R | >8 | >8 | aac(6')-Ib, aadA1, aadA2, catA1, dfrA12, EMRD, KDEA, **KPC-3**, Omp35, OmpK35, oqxB, OXA-9, SHV-11, sul1, TEM-1A | SAMN04014845 |
| 3 | AR_0005 | R | >8 | >8 | aac(6')-Ib, aph(3')-Ia, catA1, dfrA12, EMRD, KDEA, **KPC-2**, mph(A), Omp35, OmpK35, oqxA, oqxB, OXA-9, SHV-11, sul1, TEM-1A | SAMN04014846 |
| *4 | AR_0010 | S | 0.5 | 1 | CMY-94, EMRD, fosA, KDEA, Omp35, OmpK35, oqxA, oqxB14, SHV-1 | SAMN04014851 |
| 5 | AR_0016 | S | <=0.5 | <=0.5 | LEN16 | SAMN04014857 |
| *6 | AR_0034 | R | 4 | 2 | aac(3)-IId, aac(6')-Ib-G, catB3, EMRD, **IMP-4**, KDEA, mph(A), oqxA, oqxB25, qacEdelta1, qacG2, QnrB2, SHV-11, sul1, TEM-1 | SAMN04014875 |
| *7 | AR_0120 | R | 4 | > 8 | aac(6')-33, aac(6')-Ib, aadA2, aadB, aph(3')-Ia, dfrA12, KPC-2, mph(A), OmpK35, oqxA, oqxA, oqxB, sul1, sul2, TEM-1D | SAMN04014961 |
| *8 | AR_139 | I | 2 | >8 | aac(6')-IIa, armA, ARR-3, cmlA1, CMY-4, CTX-M-15, dfrA1, fosA, mph(E), msr(E), **NDM-1**, oqxA, oqxB, OXA-10, SHV-11, strA, strB, sul1, sul2 | SAMN04014980 |
| *9 | AR_0140 | S | <=0.5 | 2 | aac(3)-IId, aac(6')Ib-cr, aadA2, armA, catA1, CTX-M-15, dfrA12, dfrA14, OmpK35, OXA-181, SHV-26, sul1, sul2, tet(A) | SAMN04014981 |
| *10 | AR_0142 | S | <=0.5 | 2 | aac(3)-IId, aac(6')Ib-cr, aadA2, armA, CTX-M-15, dfrA12, dfrA14, OmpK35, OXA-181, SHV-26, sul1, sul2, tet(A) | SAMN04014983 |
| 11 | AR_0160 | R | >8 | > 8 | fosA, oqxA, oqxA, oqxB, OXA-48, SHV-11 | SAMN04015001 |
| 12 | AR_0542 | I | 2 | 2 | aac(6')-Ib, aadA2, catA1, dfrA12, EMRD, KDEA, mph(A), oqxA, oqxB, SHV-12, sul1 | SAMN05170244 |
| 13 | AR_0548 | R | >8 | >8 | aph(3'')-Ib, aph(6)-Id, CTX-M-15, dfrA14, EMRD, fosA, KDEA, **KPC-3**, oqxA, oqxB20, QnrS1, SHV-28, sul2, TEM-1 | SAMN05170307 |
| 14 | AR_0552 | R | >8 | >8 | aac(6')-Ib, aadA5, aph(3'')-Ib, aph(6)-Id, dfrA17, EMRD, KDEA, **KPC-2**, mph(A), oqxA, oqxB, SHV-11, sul1, sul2, TEM-1, tet(A) | SAMN05170144 |
| 15 | AR_0555 | R | >8 | >8 | aac(6')-Ib-G, aadA1, aph(3')-Ia, ARR-2, ble-MBL, catB, CTX-M-15, dfrA14, EMRD, KDEA, mph(A), **NDM-5**, oqxA, oqxB25, **OXA-232**, OXA-9, QnrS1, rmtF1, SHV-12, sul1, TEM-1A | SAMN11953987 |
| 16 | AR_0556 | R | >8 | >8 | aac(3)-IIa, aac(6')-Ib, aadA1, CTX-M-15, EMRD, KDEA, oqxA, oqxB20, OXA-48, OXA-9, SHV-100, TEM-1A, tet(D) | SAMN11953988 |
| *17 | AR_0558 | S | 1 | 2 | aac(3)-IId, aac(6')-Ib-cr, aadA1, aadA2, armA, ARR-2, catA1, CTX-M-15, dfrA12, dfrA14, EMRD, | SAMN11953990 |

| | | | | | | |
|---|---|---|---|---|---|---|
| | | | | | ere(A), fosA5, KDEA, OXA-181, SHV-26, sul1, sul2, tet(A), tet(R) | |
| **ATCC** | | | | | | |
| *18 | BAA-1705 | S | 1 | >8 | *blaKPC* | ATCC Download |
| *19 | 700603 | S | <=0.5 | <=0.25 | SHV-18 | ATCC Download |
| **Palo Alto Veterans Affairs Hospital** | | | | | | |
| 20 | VA1 | S | <=0.5 | <=0.5 | UNKWN | NA |
| 21 | VA2 | S | <=0.5 | <=0.5 | UNKWN | NA |
| **Johns Hopkins University** | | | | | | |
| 22 | KPU2 | S | <=0.5 | <=0.5 | UNKWN | NA |
| 23 | KPU3 | S | <=0.5 | <=0.5 | UNKWN | NA |
| 24 | KPU4 | S | <=0.5 | <=0.5 | UNKWN | NA |
| 25 | KPU5 | S | <=0.5 | <=0.5 | UNKWN | NA |
| 26 | KPU7 | S | <=0.5 | <=0.5 | UNKWN | NA |

**Supplementary Table 2.** Size shape and size parameters generated by MicrobeJ

| No. | Parameters |
|---|---|
| 1 | angularity |
| 2 | area |
| 3 | aspect ratio |
| 4 | circularity |
| 5 | curvature |
| 6 | feret |
| 7 | feret.max |
| 8 | feret.min |
| 9 | length |
| 10 | morphology |
| 11 | orientation |
| 12 | perimeter |
| 13 | pole |
| 14 | roundness |
| 15 | sinuosity |
| 16 | solidity |
| 17 | width |
| 18 | width.mean |
| 19 | width.stdev |
| 20 | trajectory |
| 21 | zscore |

**Supplementary Table 3.** Performance of various modeling strategies in classifying the strains at the first time point.

| Modeling strategy | Median Accuracy (25 resamples) |
|---|---|
| Random Forest | 0.96 |
| Naive Bayes | 0.78 |
| K-Nearest Neighbors | 0.76 |
| Support Vector Machines | 0.73 |

**Supplementary Table 4.** Performance of various modeling strategies in predicting the MIC of the strains at the first time point

| Modeling strategy | Median RMSE (25 resamples) | Median $R^2$ (25 resamples) |
|---|---|---|
| Linear Regression | 3.24 | 0.15 |
| Neural Net | 5.47 | 0.04 |
| Random Forest | 1.41 | 0.84 |
| K-Nearest Neighbors | 3.19 | 0.22 |
| Support Vector Machines | 3.92 | 0.09 |
| Gaussian Process Regression | 3.25 | 0.15 |

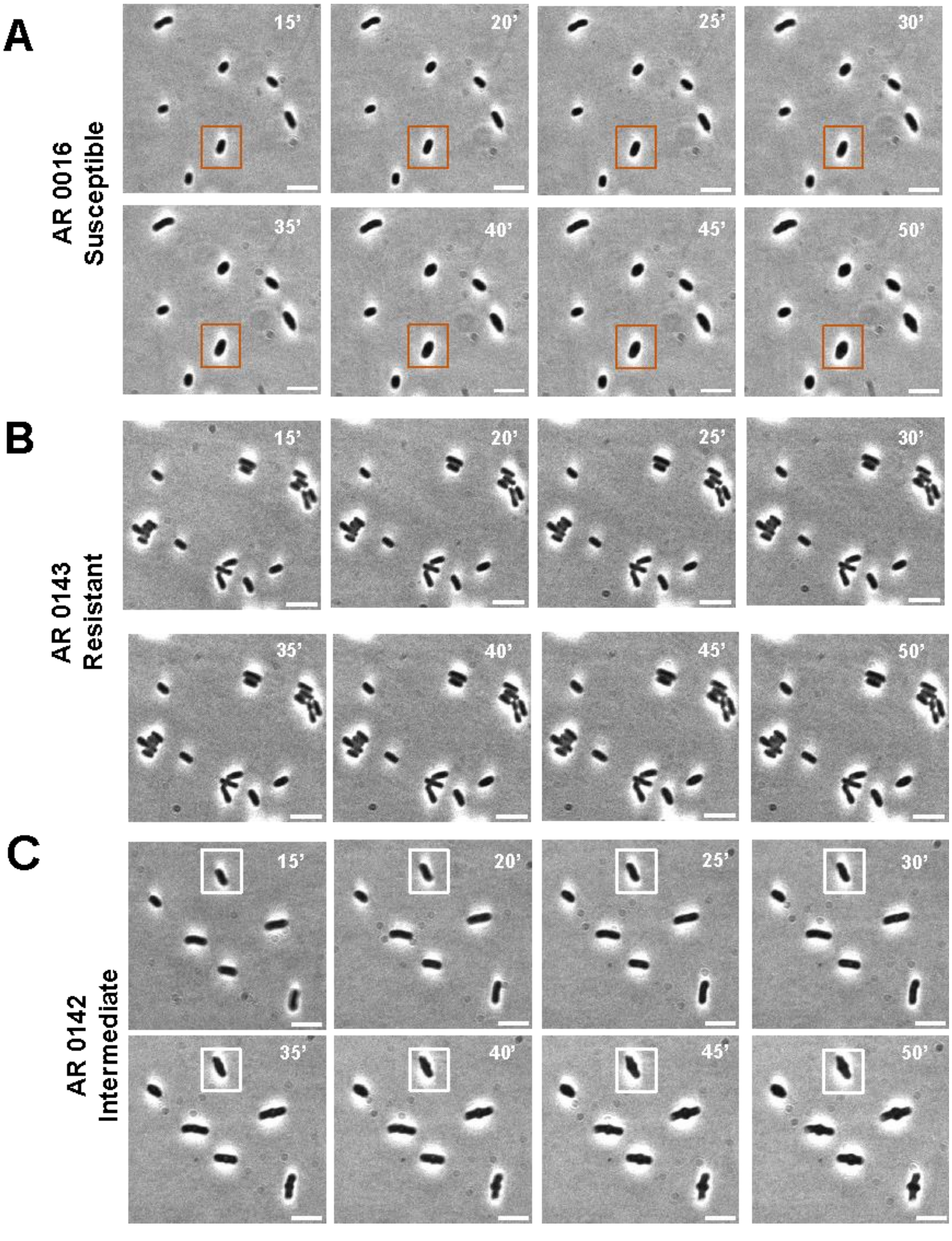


**Supplementary Figure 1.** (A-C) Morphological changes across 3 strains (KP0016 –S, KP0143 –I, and KP0142 –R) at the bacterial sub-doubling time (beginning at 15 min. and ending at 50 min.). At 5 mg/mL meropenem concentration, above the breakpoint MIC for meropenem, susceptible (orange box) and intermediate (white box) strains showed noticeable 'bulging' or protrusion around the center of the cell that was not observed in the resistant strain. Scale bars: 5 μm

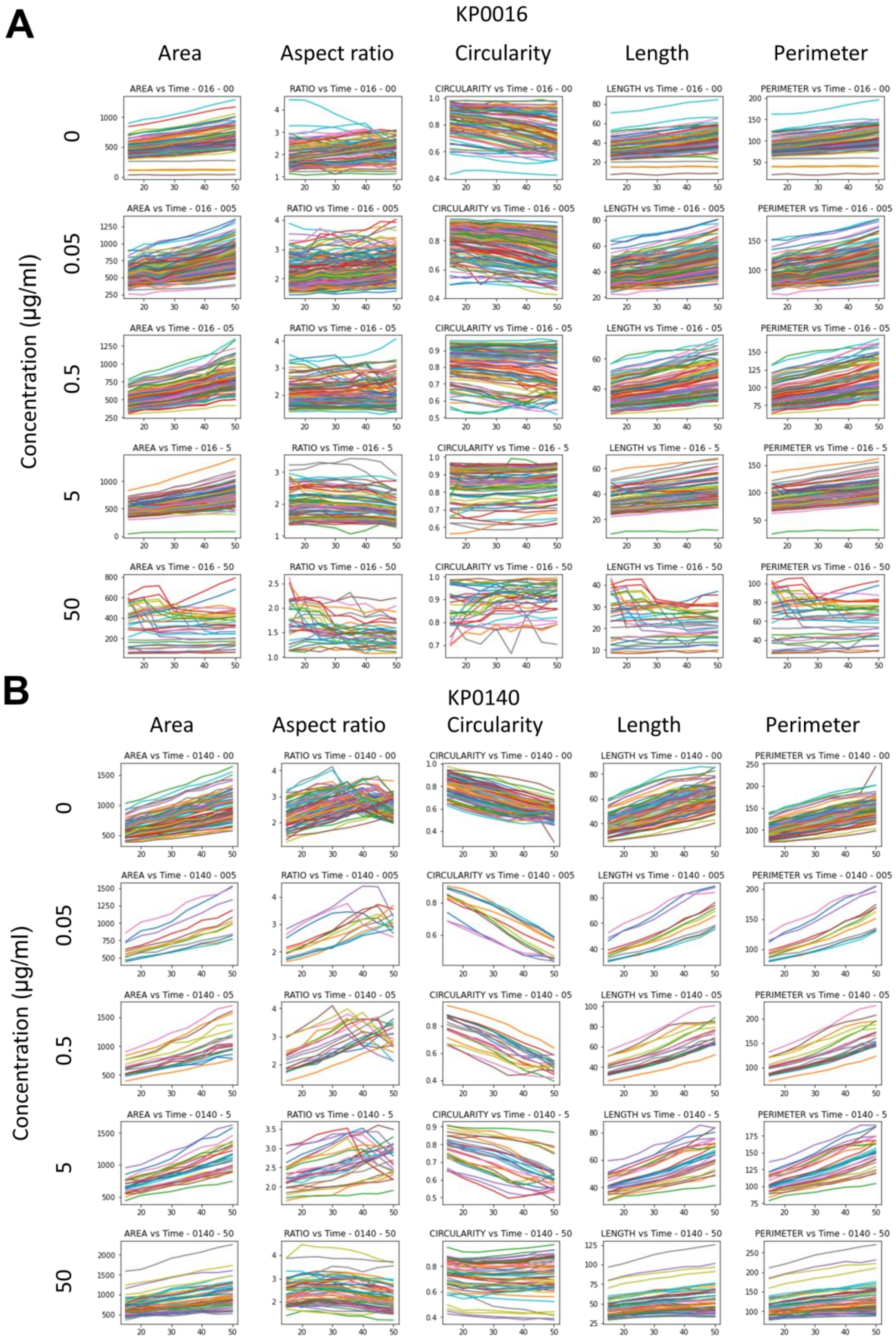


**Supplementary Figure 2.** (A-B) Dynamic morphological features of KP0016 and KP0140 from 15 min to 50 min after meropenem exposure. The experimental MICs of KP0016 and KP0140 are <= 0.5 μg/ml.

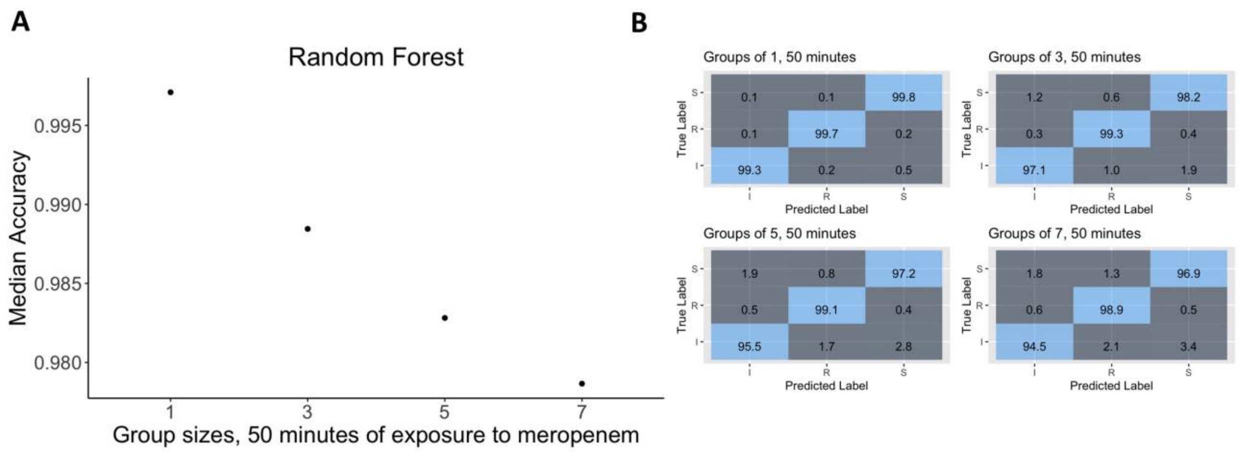


**Supplementary Figure 3.** (A) Accuracy of the random forest classifier was not improved with an increased number of cells in groups. Within 50 minutes, modeling on individual cells resulted in >99.5% accuracy but this reduces to <98% with groups of 7 cells. (B) Confusion matrices for groups of 1, 3, 5, and 7 cells after 50 minutes of antibiotic exposure.
